# DisGeNET-RDF: harnessing the innovative power of the Semantic Web to explore the genetic basis of diseases

**DOI:** 10.1101/032961

**Authors:** Núria Queralt Rosinach, Janet Piñero, Àlex Bravo Serrano, Ferran Sanz, Laura I. Furlong

## Abstract

**Motivation:** DisGeNET-RDF makes available knowledge on the genetic basis of human diseases in the Semantic Web (SW). Gene-disease associations (GDAs) and their provenance metadata are published as human-readable and machine-processable web resources. The information on GDAs included in DisGeNET-RDF is interlinked to other biomedical databases to support the development of bioinformatics approaches for translational research through evidence-based exploitation of a rich and fully interconnected Linked Open Data (LOD).

**Availability:** http://rdf.disgenet.org/

## 1 INTRODUCTION

Biomedical advancements in experimental technologies give an unprecedented capacity of description of a patient from a molecular point of view. Translational bioinformatics envisions to push forward biomedical discoveries and to enhance healthcare practice by bridging the gap between these two worlds [1]. To create this synergic translation between molecular data and clinical events, the exploration and the understanding of the complex relationships between genotype, phenotype and environment that underlie human diseases is key. DisGeNET offers one of the most comprehensive databases on gene-disease associations (GDAs) for the study of the molecular mechanisms underpinning human diseases [2]. It integrates knowledge on more than 400,000 GDAs (version 3.0) collected from a variety of authoritative sources on human genetics data and from the scientific literature. Each GDA is explicitly annotated with its supporting evidence, which makes DisGeNET a resource of reference for evidence-based knowledge discovery. Current biomedical and translational research requires leveraging and linking different types of information resources such as genetic basis of diseases, disease biomarkers, drug therapeutic applications and side effects, or effects of exposure to environmental factors. But this integration is challenging as these resources are dispersed and often technology or domain-specific. The emerging Semantic Web is gaining momentum in life sciences as it provides standards to set a semantic and syntactic interoperable infrastructure for data integration over the Web. The increasing publication of open biomedical databases structured and interlinked using the W3C Resource Description Framework (RDF) and Web Ontology Language (OWL) technologies through projects such as Bio2RDF [3] and the EBI RDF platform [4] paves the way to answer more complex and sophisticated cross-domain questions. In this paper we present DisGeNET-RDF as a new resource in the Semantic Web to foster the development of bioinformatic tools to leverage biomedical Big Data, and to facilitate knowledge navigation and discovery to support translational research.

### 2 DisGeNET-RDF DATA MODEL

DisGeNET-RDF is a linked dataset that represents GDAs as entities. These entities are semantically harmonized using the DisGeNET ontology, which formally defines types of associations between a gene and a disease in biomedical databases. The DisGeNET ontology is integrated into the Semanticscience Integrated Ontology (SIO) [5] that importantly provides ontology support for Bio2RDF Linked Data among other projects. Each GDA is made available as unique digital objects identified by Uniform Resource Identifiers (URIs), a Web-based global identification system used in RDF, and is described by different properties such as the annotated gene and disease, supporting article, SNP, and the DisGeNET score [2]. We implemented a harmonized Internationalized Resource Identifier (IRI) identification scheme for DisGeNET GDAs based on the http://rdf.disgenet.org/ domain and a unique identifier built on unique association attributes. These minted URIs are dereferenceable (it is possible to get a representation about the referenced resource on the Web), and support content negotiation for both human HTML and machine-processable RDF views. The DisGeNET-RDF data model makes extensive reuse of standard identifiers, common vocabularies and ontologies, which include OWL ontologies like the NCI thesaurus for medical vocabulary and SIO for general science. Moreover, all resources are identified by dereferenceable URIs. These URIs are from primary data providers when possible, otherwise the Identifiers.org registry of scientific identifiers is used [6]. The description of the RDF schema, ontologies used and the IRI scheme for the normalization of GDAs identification are available at the DisGeNET-RDF web site. Besides, in the context of Big Data integration and large-scale analysis over the Web, discoverability, reliability and reproducibility are major concerns for linked datasets. To address this issue, DisGeNET-RDF is available with a full provenance dataset description conformant to the W3C HCLS specification (http://www.w3.org/TR/hcls-dataset/) in order to ease its discoverability and data reuse. The W3C recommended Vocabulary of Interlinked Datasets (VoID) is used for describing the metadata of the DisGeNET-RDF dataset. In addition, it is interlinked to LOD implementations of biomedical databases and several disease terminologies such as MeSH, OMIM, Orphanet or ICD9CM available through Linked Data projects such as Bio2RDF (see statistics at http://datahub.io/ca/dataset/disgenet). Consequently, DisGeNET appears in the LOD cloud diagram (2014–08–30 version).

### 3 IMPLEMENTATION AND AVAILABILITY

DisGeNET-RDF is an open access resource of machine-processable GDAs published on the Web as Linked Data supported by a full provenance dataset description. It is created by the data providers following the Open PHACTS guidelines for exposing data as RDF (http://www.openphacts.org/). The current implementation of DisGeNET-RDF (v3.0.0) consists of 21,730,060 triples and is accessible as an RDF dump serialized in Turtle syntax for download. This dataset is made available under the Open Database License terms. The RDF dump is generated using a production system based on the D2RQ platform (http://d2rq.org/). We have also implemented a Faceted browser and a SPARQL endpoint that makes our RDF available for Linked Data navigation, information retrieval and, importantly, federated interrogation by external resources. These services are supported by Triple Store technology and they are provided by an instance of the open source edition of the OpenLink Virtuoso server (http://virtuoso.openlinksw.com/). Noticeably, we provide a web site (http://rdf.disgenet.org/) for supporting users with documentation, SPARQL query examples on how to retrieve integrated data, and contact details. In recognition of the interest in the RDF representation of DisGeNET, we note that from October 31st, 2014 to October 31st, 2015 the dataset web site had more than 11,000 page views according to Google Analytics report. Remarkably, DisGeNET-RDF is integrated in the Open PHACTS Discovery Platform [7], which is an application for drug discovery based on Semantic Web technology and RDF linked datasets.

## 4 APPLICATIONS

To identify the biological mechanisms responsible for disease etiology, pharmacological treatment and toxicological events we need to exploit biomedical data integrated in a multifaceted way. The possible applications of DisGeNET-RDF are numerous and diverse. Our SPARQL endpoint allows query federation that enables to interrogate DisGeNET with several linked open data resources with a single query. These include data on gene expression, drugs and other chemicals, biological pathways and networks, kinetic models, to just mention some of the information covered. Examples of queries are available on the Web.

## 5 CONCLUSION

DisGeNET-RDF is a new resource to harness the Semantic Web for new discovery opportunities on the genetic basis of human diseases. The publication of DisGeNET-RDF and the implementation of an SPARQL endpoint offer the possibility to integrate DisGeNET with other LOD resources to answer complex biomedical questions. DisGeNET-RDF web site supplies supporting documentation and query examples to help users getting started. Our aim is to make DisGeNET information more discoverable and to integrate it with the current open life science knowledge in order to support projects on the etiology of human diseases, drug discovery and toxicological research.

## 6 AKCNOWLEDGEMENTS

The authors thank the Open PHACTS partners, Michel Dumontier and the OpenLink staff for their input, collaboration and help. Funding: We received support from ISCIII-FEDER (PI13/00082, CP10/00524), from the IMI-JU under grants agreements n^o^ 115002 (eTOX), n^o^ 115191 (Open PHACTS)], n^o^ 115372 (EMIF) and n^o^ 115735 (iPiE), resources of which are composed of financial contribution from the European Union’s Seventh Framework Programme (FP7/2007–2013) and EFPIA companies’ in kind contribution, and the EU H2020 Programme 2014–2020 under grant agreements no. 634143 (MedBioinformatics) and no. 676559 (Elixir-Excelerate). The Research Programme on Biomedical Informatics (GRIB) is a node of the Spanish National Institute of Bioinformatics (INB).

## References

[1] R. B. Altman, “Translational bioinformatics: linking the molecular world to the clinical world.,” Clinical pharmacology and therapeutics, vol. 91, pp. 994–1000, 2012.

[2] J. Piñero, N. Queralt-Rosinach, A. Bravo, J. Deu-Pons, A. Bauer-Mehren, M. Baron, F. Sanz, and L. I. Furlong, “DisGeNET: a discovery platform for the dynamical exploration of human diseases and their genes,” Database, vol. 2015, no. 0, pp. bav028-bav028, 2015.

[3] F. Belleau, M. A. Nolin, N. Tourigny, P. Rigault, and J. Morissette, “Bio2RDF: Towards a mashup to build bioinformatics knowledge systems,” Journal of Biomedical Informatics, vol. 41, no. 5, pp. 706–716, 2008.

[4] S. Jupp, J. Malone, J. Bolleman, M. Brandizi, M. Davies, L. Garcia, A. Gaulton, S. Gehant, C. Laibe, N. Redaschi, S. M. Wimalaratne, M. Martin, N. Le Novère, H. Parkinson, E. Birney, and A. M. Jenkinson, “The EBI RDF platform: Linked open data for the life sciences,” Bioinformatics, vol. 30, no. 9, pp. 1338–1339, 2014.

[5] M. Dumontier, C. J. Baker, J. Baran, A. Callahan, L. Chepelev, J. Cruz-Toledo, N. R. Del Rio, G. Duck, L. I. Furlong, N. Keath, D. Klassen, J. P. McCusker, N. Queralt-Rosinach, M. Samwald, N. Villanueva-Rosales, M. D. Wilkinson, and R. Hoehndorf, “The Semanticscience Integrated Ontology (SIO) for biomedical research and knowledge discovery.,” Journal of biomedical semantics, vol. 5, no. 1, p. 14, 2014.

[6] N. Juty, N. Le Novère, and C. Laibe, “Identifiers.org and MIRIAM Registry: community resources to provide persistent identification.,” Nucleic acids research, vol. 40, no. Database issue, pp. D580–6, 2012.

[7] A. J. Gray, P. Groth, A. Loizou, S. Askjaer, C. Brenninkmeijer, K. Burger, C. Chichester, C. T. Evelo, C. Goble, L. Harland, S. Pettifer, M. Thompson, A. Waagmeester, and A. J. Williams, “Applying linked data approaches to pharmacology: Architectural decisions and implementation,” Semantic Web, vol. 5, no. 2, pp. 101–113, 2014

